# A rich get richer effect governs intracellular condensate size distributions

**DOI:** 10.1101/2022.03.08.483545

**Authors:** Daniel S.W. Lee, Chang-Hyun Choi, David W. Sanders, Lien Beckers, Joshua A. Riback, Clifford P. Brangwynne, Ned S. Wingreen

## Abstract

Phase separation of biomolecules into condensates has emerged as a ubiquitous mechanism for intracellular organization and impacts many intracellular processes, including reaction pathways through clustering of enzymes and their intermediates. Precise and rapid spatiotemporal control of reactions by condensates requires tuning of their sizes. However, the physical processes that govern the distribution of condensate sizes remain unclear. Here, we utilize a combination of synthetic and native condensates to probe the underlying physical mechanisms determining condensate size. We find that both native nuclear speckles and FUS condensates formed with the synthetic Corelet system obey an exponential size distribution, which can be recapitulated in Monte Carlo simulations of fast nucleation followed by coalescence. By contrast, pathological aggregation of cytoplasmic Huntingtin polyQ protein exhibits a power-law size distribution, with an exponent of −1.41 ± 0.02. These distinct behaviors reflect the relative importance of nucleation and coalescence kinetics: introducing continuous condensate nucleation into the Monte Carlo coarsening simulations gives rise to polyQ-like power-law behavior. We demonstrate that the emergence of power-law distributions under continuous nucleation reflects a “rich get richer” effect, whose extent may play a general role in the determination of condensate size distributions.

## Introduction

Condensates of biological macromolecules play critical roles in many biological processes, including ribosome synthesis^1^, DNA organization^2^ and repair^3^, and stress responses^4,5^. In the context of metabolism, co-clustering enzymes into condensates can increase the efficiency of reactions by spatially coordinating intermediates^6^, but only for a restricted range of sizes. Similarly, large deviations from typical condensate sizes, which have been described for many nuclear condensates, including nucleoli^1^, Cajal bodies^7^, and nuclear speckles^8^, are associated with dysfunction and pathology. For instance, increased nucleolar size is associated with increased ribosome biogenesis^9^ and has been associated with both cancer^10^ and Hutchinson-Gilford progeria^11^. Moreover, condensates are strongly linked to pathological aggregation diseases such as Alzheimer’s and ALS. In these cases, there are many remaining questions about how large condensates might nucleate irreversible aggregates, and whether large aggregates are themselves pathological, or merely end-stage outcomes from smaller pathological assemblies.

In non-living, colloidal systems, the kinetics of coarsening processes are well-known to be inherently linked with and to generally determine cluster-size distributions^12–15^. By contrast, while biomolecular condensates have been observed in living cells for many years, their size distributions and coarsening dynamics have rarely been examined in detail. Among the few previously studied examples are nucleoli in *Xenopus laevis* and in cultured human cells, which have been shown to have power-law size distributions^1,16^. Interestingly, such broad, scale-free size distributions are at odds with the suggestion that endogenous condensates have well-defined, functionally regulated sizes. However, in more typical rapidly dividing mammalian cells, nuclear bodies, such as nucleoli and speckles^8,17^, tend to appear following mitosis and coalesce^18^, but then maintain relatively stable sizes until the next cell division, likely in part due to strong mechanical constraints from chromatin^19–22^. This picture is consistent with other recent work that has taken advantage of synthetic nuclear condensates, which form quickly^23^ but grow and coalesce slowly due to their constrained, subdiffusive motion^20^.

In some pathological contexts, cytoplasmic aggregates known as inclusion bodies or aggresomes are continuously produced, potentially providing a continually changing size distribution. For example, in the case of Huntington’s disease, the progressive misfolding of mutant Huntingtin protein exhibiting abnormally large stretches of polyglutamine or “polyQ” repeats leads to steady accumulation of material into irreversible aggregates. Although generally more solid-like than most liquid-like physiological condensates, pathological aggregates can also generally move throughout the cell, and stick or fuse upon collision, forming progressively larger structures that may even span the entire cell^24^ and are associated with neuronal dysfunction^25^ and cytotoxicity. How such steady accumulation of aggregating material is similar to or differs from the coarsening dynamics of physiological condensates, and the physical and biological principles that underlie such phenomena more generally, remain poorly understood.

Here, we combine live-cell experiments and simulations to elucidate general principles linking condensate growth dynamics and size distributions. We demonstrate that endogenous nuclear speckles, which exhibit fast nucleation followed by gradual coalescence, yield an exponential size distribution, which can also be recapitulated in an engineered intracellular phase-separating system. By contrast, cytoplasmic Huntington aggregates show that continuous material production instead leads to a power-law distribution. We describe the differences that account for the scaling forms of the condensate size distribution in terms of a “rich get richer” effect, wherein growth by merger is biased towards larger condensates. These findings may provide insight into how cells biophysically regulate the size and number of condensates.

## Results

### Endogenous and synthetic nuclear condensates demonstrate an exponential size distribution

We first sought to determine the distribution of the sizes of an endogenous nuclear condensate, the nuclear speckle. Previous work suggested that speckles form quickly after mitosis and grow by coalescence^8^, thus acting as a model for the dynamics of a typical condensate. We imaged stem cells (iPSCs) in which SON, a known marker of speckles^26^, was endogenously tagged with GFP (Fig. 1a), since overexpressing condensate constituents can alter condensate size^27^. We then segmented the speckles in 3D and quantified the probability density *f*(*V,t*) of condensate volume *V*at time *t*. Because the size distribution is highly sensitive to binning and cell-to-cell variation, we first considered the cumulative distribution function (CDF) of condensate sizes for each cell:

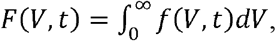

and rescaled these distributions by each cell’s average condensate volume. We averaged over the rescaled distributions to account for cell-to-cell variation and finite sampling, and plotted the complementary cumulative distribution (CCDF), CCDF = 1 −*F*. We found that the nuclear speckle CCDF as a function of condensate volume *V* is linear on a semi-log plot for all timepoints, implying that these condensates are well-described by an exponentially decaying size distribution (dashed line, Fig. 1b)

**Fig. 1:**
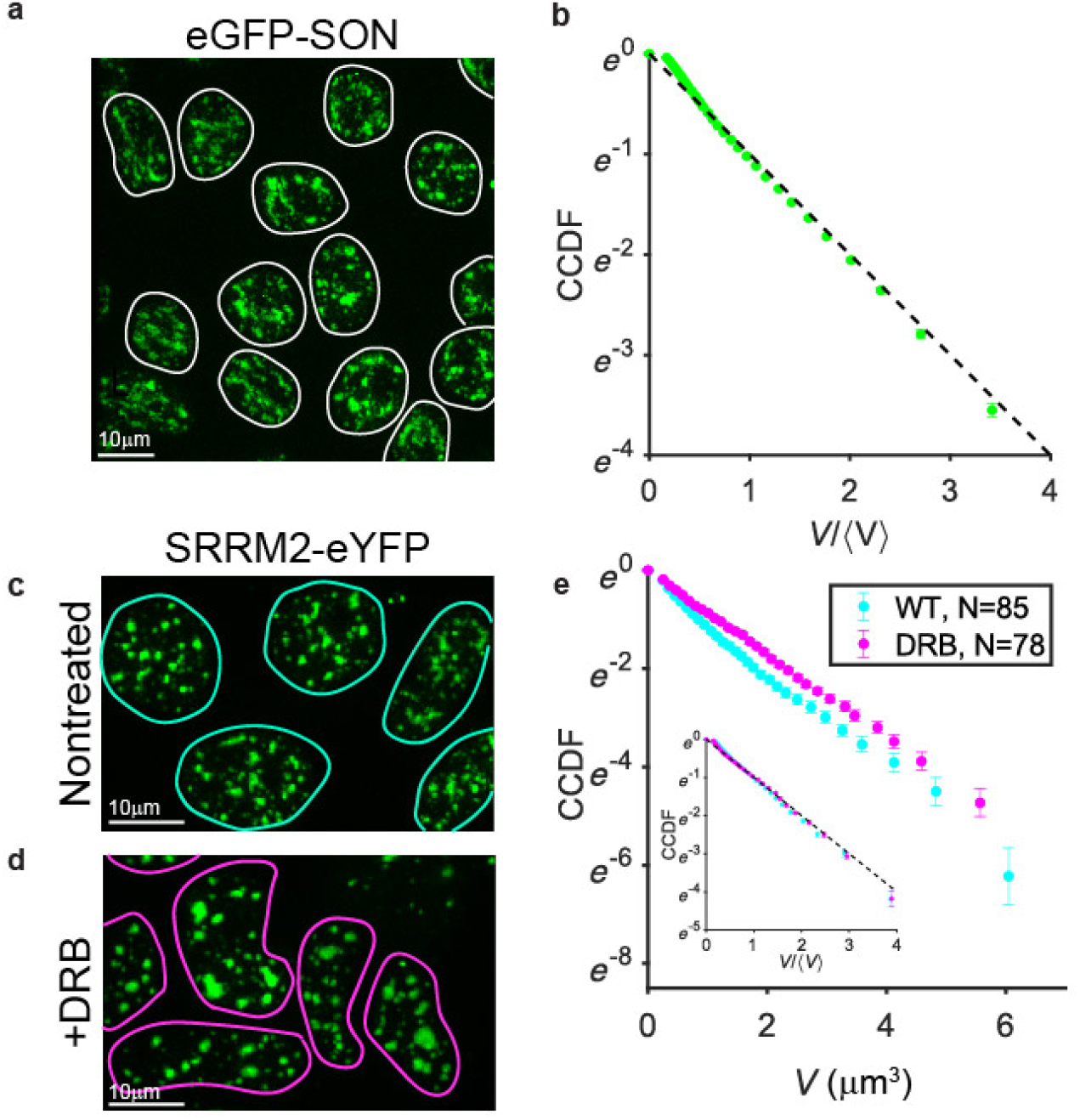
Endogenous nuclear speckles display exponential distributions before and after transcriptional inhibition. a. Example maximum-projected image of induced pluripotent stem cells tagged with eGFP-SON and imaged in three dimensions. Nuclear outlines are highlighted in white. b. For each nucleus (*N* = 453), speckles were segmented in 3 dimensions, and the complementary cumulative distribution function (CCDF) calculated, rescaled, and averaged, revealing good agreement with an exponential distribution (black dashed line). Error bars reflect standard error of the mean over cells. c. Nuclear speckles were labeled in HEK-D cells by tagging SRRM-2 with eYFP and imaged in 3 dimensions; nuclear outlines highlighted in cyan. d. Treatment with the transcriptional inhibitor DRB resulted in larger, brighter speckles and aberrant nuclear morphology. Nuclear outlines are highlighted in magenta. e. The CCDF was calculated for both nontreated control (*N* = 85; cyan points) and following 4 hours of treatment with 50μg/mL DRB (*N* = 78; magenta points), resulting in an increase in average speckle size from 0.92 ± 0.02 μm^3^ to 1.16 ± 0.03 μm^3^(mean±SEM), giving linear CCDFs with slightly different slopes on a semi-log plot. Collapsing by rescaling by average speckle volume (inset) confirms that both distributions remain exponential. Error bars reflect standard error of the mean over cells.

To further interrogate this behavior in a more tractable cell line, we utilized a monoclonal HEK293-derived (“HEK-D”; see Methods) cell line with eYFP-tagged SRRM2, another speckle marker (Fig. 1c)^26^, and expressing mCh-tagged NPM1 to mark nucleoli. We first verified that an exponential was still observed in this cell line (Fig. 1e, dashed black line). Previous work^8^ suggested that treatment with the transcriptional inhibitor 5,6-Dichlorobenzimidazole 1-β-D-ribofuranoside (DRB) accelerates long-distance collisions between speckles, facilitating fusion. We asked whether this increase in dynamics would alter the form of the size distribution. Following 4 hours of treatment with DRB, we find an increase in mean speckle size from 0.92 ± 0.02 μm^3^ (mean ± standard error of the mean over 85 cells) in the nontreated control to 1.16 ± 0.03 μm^3^ (78 cells), consistent with the literature. However, despite this average size increase, both distributions were exponential (Fig. 1e). Similar results were seen using Actinomycin D (Fig. S1). By contrast, in nucleoli, we observed a power-law distribution with a slope of approximately -1 (Fig. S1)^16^, and a marked decrease in nucleolar size after both ActD and DRB treatment^28^, both consistent with previous work.

To examine the origin of these distributions in an even more experimentally tractable platform, we utilized the Corelet optogenetic system whose nucleation and coarsening dynamics have been well-characterized^20,23,29^. Briefly, the Corelet system consists of two components (Fig. 2a): a 24-mer ferritin core whose monomers are fused to a green fluorescent protein (GFP), and an iLiD (improved light-induced dimer) domain plus a protein, here the intrinsically disordered region (IDR) of FUS, fused to an mCherry (mCh) fluorophore and an sspB domain (stringent starvation protein B). Upon blue light activation, the iLiD and sspB heterodimerize, resulting in oligomerization of the FUS IDR, which drives phase separation within seconds after light activation^29^. Using this biomimetic system expressed in human osteosarcoma (U2OS) cells, we activated and imaged for 105 minutes^20^; we observed that the CCDF is again consistent with an exponentially decaying size distribution. By examining the distribution as a function of time since initiation of condensate formation with light, we find that for a particular cell, the CCDF retains the linear shape on a semi-log plot, but changes slope as the condensates coarsen and the average condensate size grows (Fig. 2b). To confirm that this behavior was consistent across cells and timepoints, we rescaled each CCDF by mean condensate size ⟨*V* (*t*) ⟩ for each cell and time, revealing that these distributions obeyed the same scaling, all collapsing onto a line with slope -1 (dashed black line) corresponding to an exponential distribution (Fig. 2c). Finally, to verify the Corelet exponential cluster-size distribution, we also calculated and compared the volume-weighted or “particle-centric” mean ⟨*V*⟩_p_ and variance *σ*^2^_p_ for each nucleus and timepoint, which should be less noisy but obey the same relation as the cluster-centric distribution for a system with fixed density clusters (e.g., LLPS). We note that the points fall along the ⟨*V*⟩_p_^2^ = *σ*^2^_p_ line expected for an exponential distribution, with a slight horizontal shift which can be explained by the bias in the mean arising from a minimum detection size (Supplementary Note; Fig. S2).

**Fig 2:**
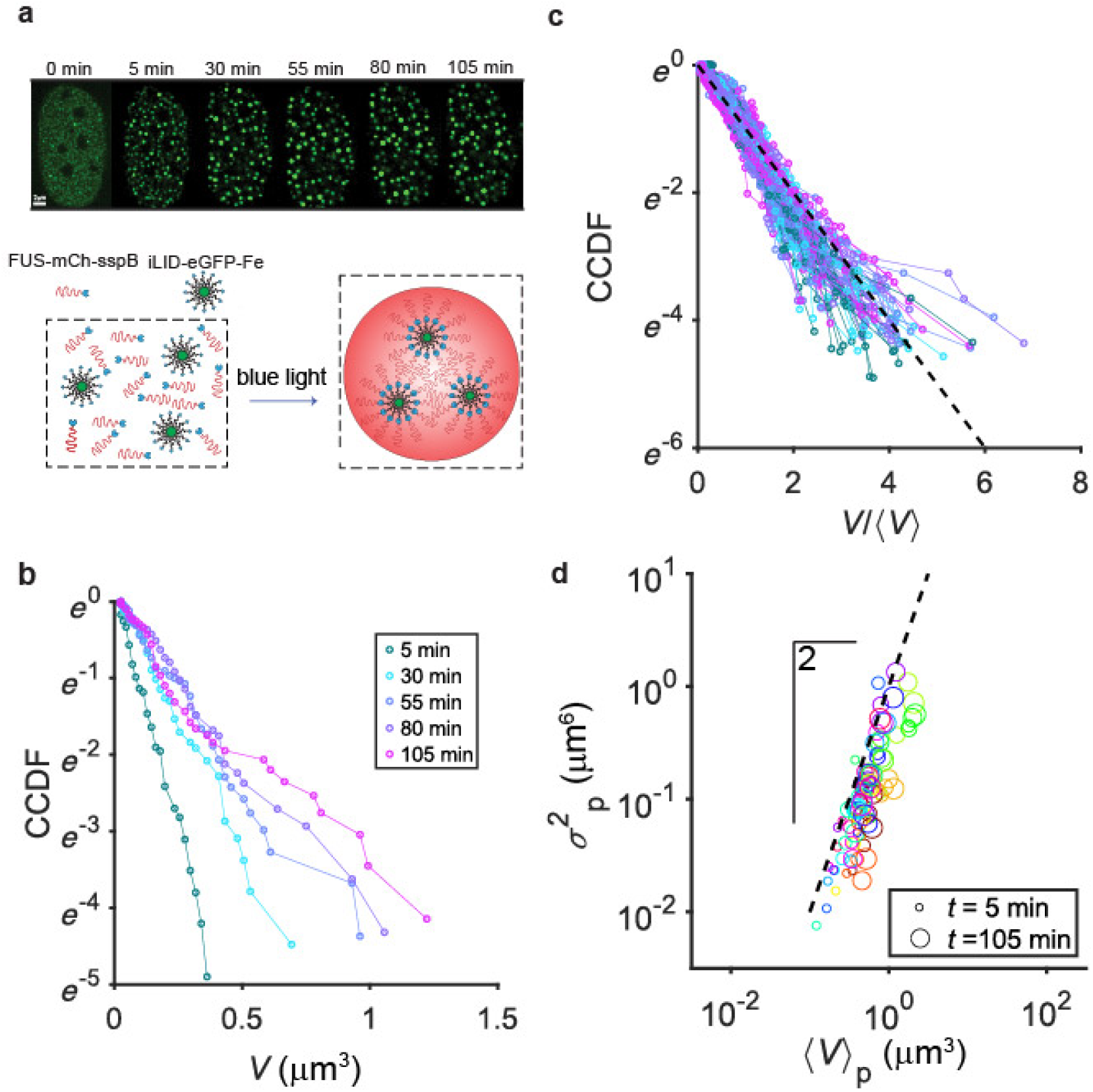
Synthetic nuclear condensates display exponential distributions as they coarsen. a. The synthetic Corelet system produces nuclear condensates on demand. The FUSN-mCherry-sspB and iLID-eGFP-ferritin fusion proteins are co-expressed in the nucleus. In the presence of blue light, sspB and iLiD bind quickly, decorating the 24-mer ferritin core with oligomerized FUSN, driving oligomerization and phase separation (top row). b. The CCDF of condensate volumes for the nucleus shown in Fig. 1A plotted semi-log reveals a progressively decreasing slope as the droplets coarsen in time following the initial quench (<5 minutes) and the mean condensate size increases. c. For 18 nuclei, CCDFs rescaled by mean condensate volume were obtained at the timepoints depicted in Fig. 1B (reanalyzed data from Lee 2021), and compared to the expectation for an exponential distribution (dashed black line). d. To further verify that the distribution was exponential, we calculated the mass-weighted or particle-wise mean and variance, defined as 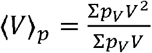 and 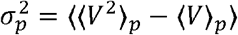 respectively. These quantities were calculated for each nucleus every 25 minutes starting from 5 minutes to 105 minutes following initial activation, variance was plotted against mean on a log-log plot, and compared to the expected slope of 2 for an exponential distribution.

### Monte Carlo simulations recapitulate exponential distributions, regardless of subdiffusion

To quantitatively understand the underlying basis for the exponential size distribution of condensates, we developed Monte Carlo simulations to replicate and manipulate the coagulation dynamics. We hypothesized that, in general, a fast quench, wherein many small condensates nucleate nearly simultaneously, followed by growth via slow coagulation, would produce an exponential distribution. In our simulations, we first generated *N* spheres whose initial volumes were sampled from a uniform distribution in the range [1,2], which were randomly placed in a cubic box of side length *L*. Spheres whose initial positions overlapped were merged by replacing them with a larger sphere centered at their center of mass and conserving volume (Fig. 3a). The spheres were then allowed to move according to fractional Brownian motion with an exponent *α*, i.e., ⟨*r*^2^⟩ = 2*dDτ*^*α*^ for displacement 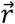, dimension *d*, diffusion coefficient *D*, and time lag *τ*. Thus, the simulations could be tuned to mimic subdiffusive (*α* < 1) or diffusive (*α* = 1) behavior, a physiologically relevant parameter, considering the apparent ubiquity of subdiffusion in cells^20,30,31^. In the simulations, spheres diffused according to the specified dynamics and were merged upon contact. We found that the size-dependent rate of mergers agreed with predictions from coagulation theory (Fig. S3)^32^.

**Fig. 3:**
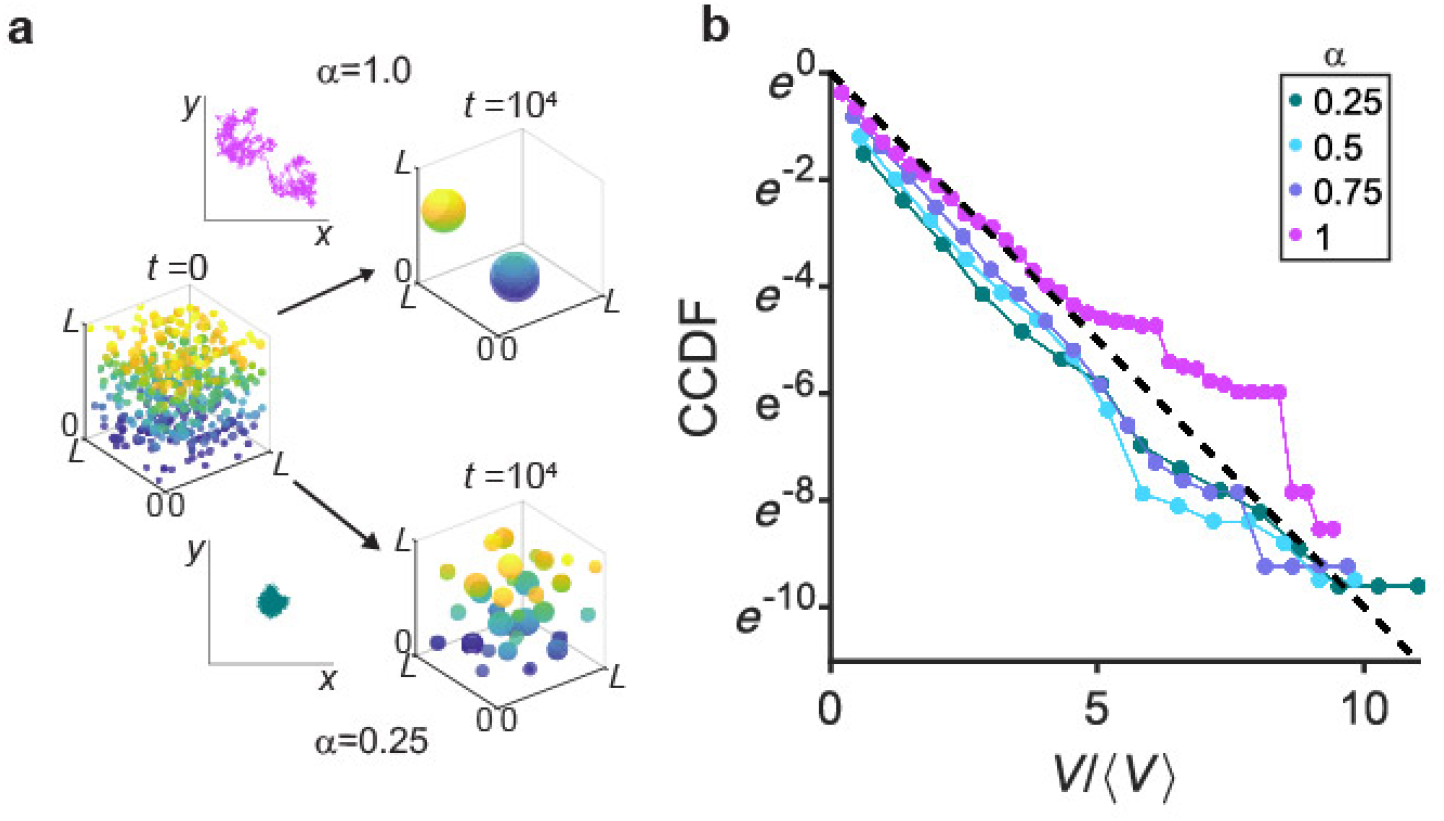
Monte Carlo simulations of subdiffusive coalescence also produce exponential distributions. a. Spheres were randomly placed in a box with periodic boundary conditions and allowed to coalesce into larger spheres upon collision. Sphere dynamics are dictated by fractal Brownian motion (*cf*. example trajectories), resulting in faster coalescence for larger subdiffusive exponent α (see Materials and Methods). b. Simulations of 1000 spheres with initial volumes drawn from a uniform distribution between 1 and 2 were run for 10^4^ timesteps at a volume fraction of 5%, for varying subdiffusive exponents α from 0.25 to 1. Distributions first were averaged over 20 replicates before calculating the CDF and CCDF. The CCDF, rescaled by mean sphere volume, was plotted semi-log for comparison to an exponential (black dashed line).

We ran replicates over a range of (sub)diffusive conditions and calculated the distribution of volumes after a fixed time. Plotting the sphere-size distribution and rescaling by the average sphere volume as in the analysis of the experimental data, we observed exponential distributions under all conditions (Fig. 3b). Thus, in simulations with a wide range of diffusive exponents and in experiments with various synthetic and endogenous condensates, we observed exponential distributions.

### Huntingtin polyQ aggregates exemplify the slow production regime and have a power-law condensate-size distribution

Our findings thus far suggest that rapid nucleation of small condensates followed by slower coarsening due to coalescence of condensates generically results in an exponential distribution. To further elucidate how the relative timescales of nucleation and coagulation affect the scaling of the condensate-size distribution, we next considered a contrasting system characterized by continuous, sustained nucleation of new small condensates. Specifically, we utilized the Huntingtin polyQ exon 1 (Htt-polyQ) system, which, unlike the above systems, exhibits a slow and steady increase in total aggregate material with time. Exogenous eGFP-Htt-polyQ protein was lentivirally expressed in HeLa cells to induce aggregate formation. Consistent with previous work^33^, eGFP-Htt-Q31 did not form aggregates, but eGFP-Htt-Q73 formed large aggregates in perinuclear regions of the cytoplasm within 4 days following infection (Fig. 4a). These aggregates were observed to nucleate over the span of hours and merge upon contact (Fig. 4b). To quantify production dynamics, we measured the total projected area of aggregated material as a function of time, finding a roughly constant average production rate of approximately 0.04 ± .01μm^2^/s (Fig. 4b), which corresponds to an increase of 2.28 ± 0.24 fold over the observed interval (Fig. S4); this contrasts with the Corelet system, which generally remained stable in total condensate volume, following an initial nucleation period (Fig. S2 a,b).

**Fig. 4:**
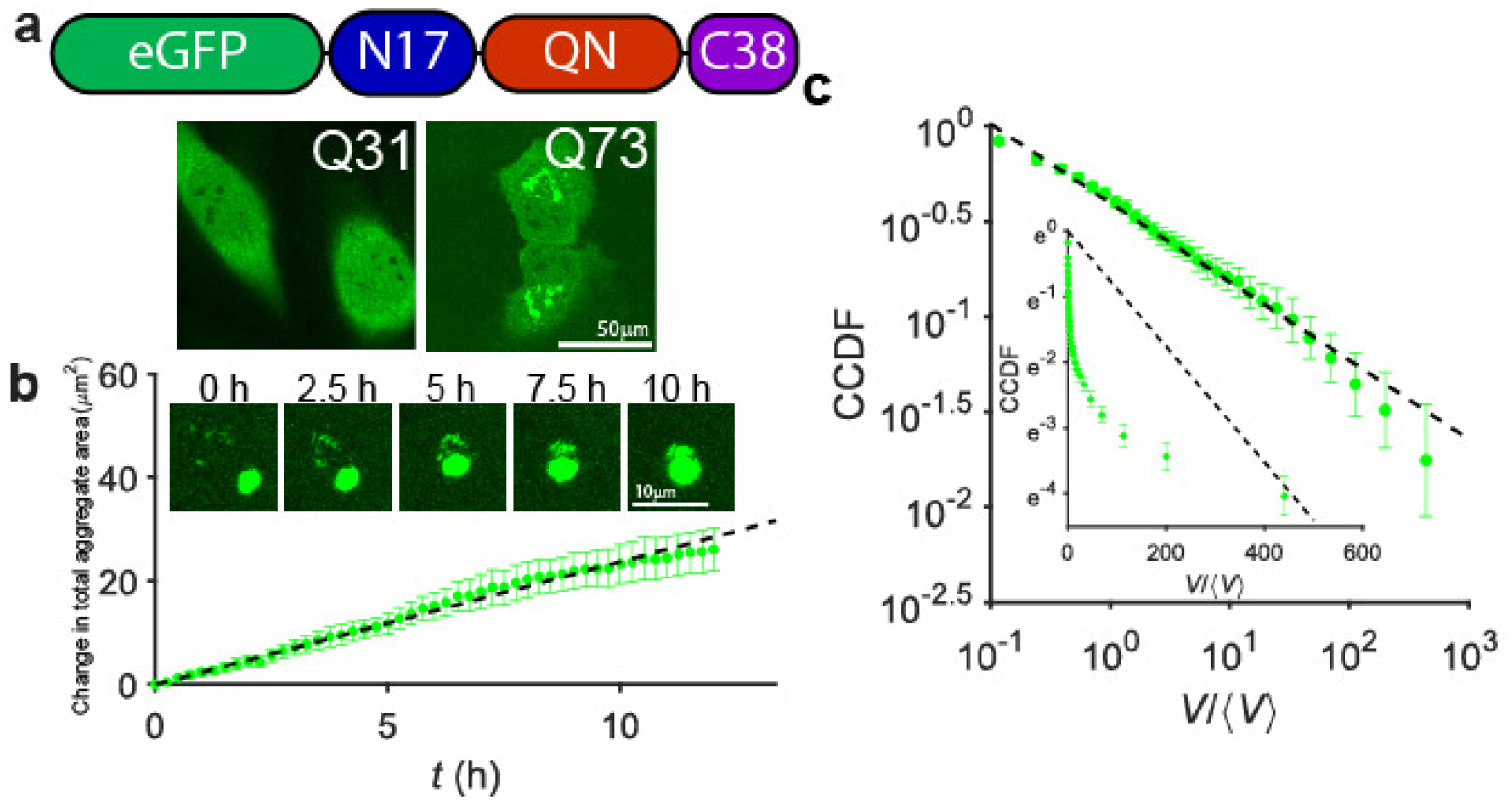
PolyQ aggregates exhibit slow nucleation dynamics and a power-law cluster-size distribution. a. HeLa cells were transduced with lentivirus to express eGFP-Htt-PolyQ constructs of mutant Huntingtin exon 1 composed of N17, QN, and C38 domains. Four days following transduction, cells infected with eGFP-Htt-Q31 had a diffuse GFP signal, while bright micron-scale puncta were evident in those infected with eGFP-Htt-Q73. b. Cells expressing eGFP-Htt-Q73 were imaged for 12 hours. Aggregates were observed to merge upon contact (inset). PolyQ growth was quantified over *N* = 5 cells imaged every 15 minutes for 12 hours. Subtracting the initial projected aggregate area and averaging revealed an increase in aggregate area over time at a rate of 0.04 ± .01 μm^2^/s. c. PolyQ aggregate cluster-size distribution was calculated by averaging per-cell cumulative distribution functions over cells (*N* = 114) between 4 and 6 days after lentiviral infection. The CCDF was fit to a power law, yielding CCDF(*V*) ∼ *V*^0.41±0.03^ . The CCDF was also compared to the expectation for an exponential distribution (inset).

To examine the effect of these contrasting kinetics, we quantified the size distributions of polyQ aggregates. To calculate the per-cell distribution and avoid binning artifacts, a CDF was calculated for each cell and averaged over cells. The resulting CCDF was well-fit by a power law with exponent −0.41 ± 0.03, corresponding to an exponent of −1.41 for the PDF (Fig 4C); the CCDF was poorly-fit by an exponential (inset).

### Slow injection results in a broad condensate-size distribution

We hypothesized that the broad power-law distribution observed in the polyQ system is due to the similar timescales for the appearance of new aggregates and mergers of existing aggregates. To test this idea, we returned to our Monte Carlo simulations to model the slow nucleation of the polyQ system. Specifically, we modified the simulations such that spheres appear at a fixed injection rate of *J* spheres per timestep, rather than all at once at *t* = 0 (Fig. 5a). For a range of *J* values, we ran simulations until a fixed equal number of spheres were injected, and then calculated the CCDFs (Fig. 5b). For relatively high values of *J* (*J* > 0.1), the resulting distributions were approximately exponential, but for low values of *J* (*J* < 0.1), the distributions were more power law-like (Fig. 5b), consistent with previous work^1,34^. In particular, we note that the *J* = .01 case produces a power law over two decades with an exponent of − 1.51 ± 0.13, very close to the power-law exponent observed in our polyQ experiments. This suggests that the slow addition of new material drives the system towards a broader distribution of sizes, associated with a power-law distribution with exponent −1.5.

**Fig. 5:**
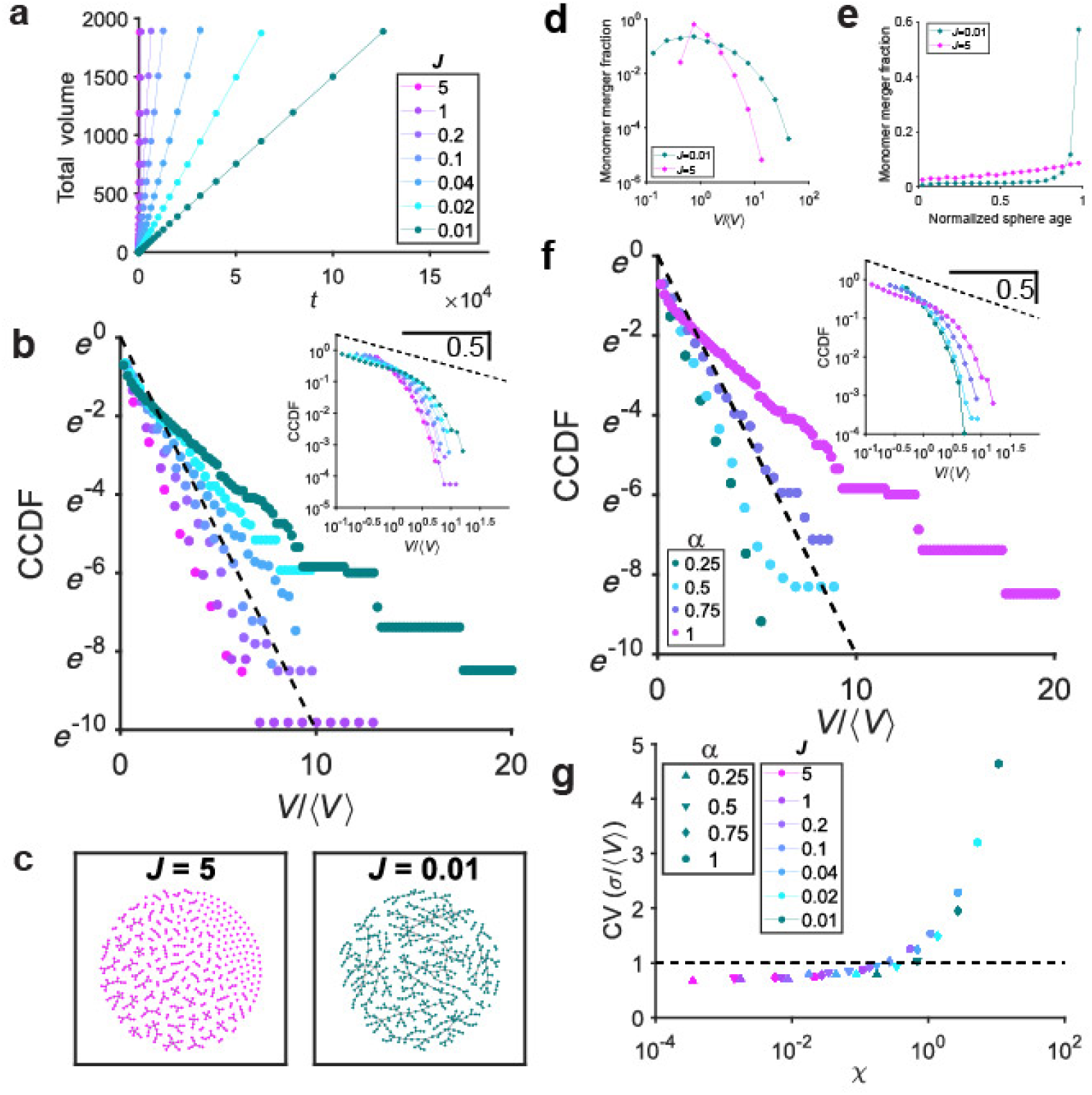
Simulations incorporating gradual material production demonstrate a power law. a. Simulations were modified to incorporate the gradual introduction of material at a rate of *J* spheres per unit time, starting with an empty simulation cell of total volume such that following injection of 1200 spheres whose initial sizes were again drawn from a uniform distribution between 1 and 2, a volume fraction of 4% was obtained. This protocol is summarized by the plot showing total particle volume increasing with time with a slope corresponding to *J*. b. The cluster-size distribution was calculated for systems following steady injection of 1200 spheres and plotted both semi-log and log-log (inset). Conditions with high *J* fit an exponential distribution, as shown in CCDF, while lower *J* gave a better fit to a power law with slope close to -0.5 based on log-log plot of CCDF (inset, dashed black line). c. Merger events between spheres were mapped for *J* = 5 (left, magenta) and *J* = .01 (right, dark green) for systems following injection of 50 spheres each. Each node represents an injected sphere; edges between nodes signify mergers during the course of the simulation. d. Mergers involving “monomer” spheres (i.e., spheres of the minimum size) were analyzed for *J* = .01 and *J* = 5. The distribution of sizes of spheres with which ‘monomeric’ spheres merged, normalized by the mean sphere size at the moment of merger was calculated and plotted for 960 simulation replicates of each condition. e. The “age” of spheres, normalized by total simulation duration (i.e., the time it took to inject 1200 spheres at varying *f*) with which monomeric spheres merged was calculated for 960 replicates of *f* = .01 and *f* = 5. f. Injection simulations were run with *f* = .01 for varying *α*. The CCDF was calculated and plotted, demonstrating that for low *α*, an exponential distribution is obtained. The CCDF for the subdiffusive injection case was also calculated and plotted for comparison to a power law (inset). g. For each *α* and *J*, simulations were run until approximately 1200 particles were injected. The coefficient of variation was calculated for the resulting distributions and plotted as a function of *χ*, the ratio of the merger timescale to the injection timescale.

Given that our simulations yielded either an exponential or a power-law distribution simply by changing the rate of injection of new material, this provided an opportunity to track down the underlying mechanism responsible for these qualitatively different outcomes *in silico*. Since the size distribution depends on which mergers occur, we mapped the collision events of the respective systems to obtain the network structure of mergers. To visualize this merger network, we depicted each sphere as a unique node in a graph representation, drawing an edge between it and another sphere to represent a merger event at some point in the simulation (Fig. 5c). In the high *J* case, we observed many isolated sets of connected nodes, consistent with the exponential size distribution, while for low *J*, the network is highly connected, yielding fewer total spheres but with a much broader range of final volumes.

For low *J*, the merger network structure is reminiscent of preferential attachment phenomena, which have been shown in a wide variety of contexts to generate power-law distributions^35–37^. In the case of low *J*, the first spheres to be injected have more opportunities to merge and consume smaller spheres, resulting in preferential attachment onto larger spheres or a “rich get richer” effect. By contrast, in the case of high *J*, the sphere merger probability is approximately independent of size.

We verified this quantitatively by analyzing mergers in the *J* = 5 and *J* = .01 cases. We considered mergers involving “monomeric” spheres, i.e., spheres which had been injected and not yet undergone any mergers. We recorded the size of the spheres with which they merged, normalized by the mean sphere size at the moment of merger, finding that for *J* = 5, monomeric spheres typically merged with spheres close to the mean sphere size, while for *J* = .01, the distribution of merger partners was much wider, reflecting the bias of collisions towards spheres much larger than the mean (Fig. 5d). Furthermore, over the course of the simulations, we calculated the age of the spheres with which monomer spheres collided, normalizing by the total duration of the simulation, finding that merger fraction increases linearly with sphere age in the fast injection case, as would be expected if merger probability is size-independent. However, in the slow injection case, the merger fraction was even more strongly dependent on the age of the merger partner, with approximately 80% of monomeric spheres merging with a sphere that had been injected in the first 25% of the simulation (Fig. 5e), matching our intuition that mergers were heavily biased towards “older” spheres.

We reasoned that if decreasing *J* results in a broader distribution because injection becomes slower relative to merger, then conversely, slowing down the merger rate should similarly recover an exponential distribution, since having a modest head start on merger opportunities would not be consequential if the merger time is very long. Therefore, we held *J* fixed at a value of 0.01 and varied the subdiffusive exponent in our simulations. We found that decreasing the subdiffusive exponent and thus slowing down mergers indeed recovered an exponential distribution (Fig. 5f).

To characterize the crossover between an exponential and a power-law size distribution, we ran simulations varying both *α* and *J*. To determine which process was kinetically limiting, we computed and compared the merger timescale *τ*_merge_ to the injection timescale *τ*_inject_ based on input parameters. The merger timescale *τ*_merge_ is given by the time required to close the distance between spheres by (sub)diffusion, i.e., 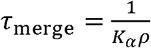, where *ρ* is total number of injected spheres per system volume, and *K*_*α*_ is the collision rate constant, whose dependence *α* is nontrivial and was calculated empirically in simulation (Fig. S4). The injection timescale *τ*_inject_ is the time required to inject all the spheres into the system, i.e., 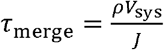. We then define the dimensionless number *χ*:

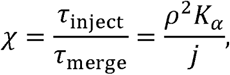

where 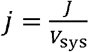 is the rate of injection of new spheres per unit volume. For each simulation condition, to characterize the distribution of sphere volumes at the time when the last new sphere was injected, we calculated the ratio of the standard deviation to the mean, aka the coefficient of variation (CV). For an exponential distribution, CV = 1, while a power law-like distribution will have a larger CV. We then plotted *χ* against the CV for each simulation condition (Fig. 5g) and found that all conditions collapsed onto a single curve, matching our intuition that the ratio of timescales captured by *χ* governs the final distribution. For *χ* < 1, the timescale of a typical merger is greater than the time required to inject all the material into the system, so the distribution remains narrow and close to an exponential, since mergers are sparse and occur between random pairs. However, for *χ* > 1, many merger events will occur during the injection process, giving “older” spheres more merger opportunities, mimicking the polyQ case, facilitating the “rich get richer” effect, and resulting in a broader, more power law-like distribution.

### The “rich get richer” effect generates power-law distributions

The “rich get richer” effect can also be understood in the context of the simplest coagulation framework first analytically solved by Smoluchowski^38^, wherein the relative rate at which spheres of particular volumes (*V*_1_, *V*_2_) with diffusion coefficients (*D*_1_, *D*_2_), collide can be described using a matrix known as a coagulation kernel *K*(*V*_1_, *V*_2_). For the diffusive spheres in our simulations, this is given by:

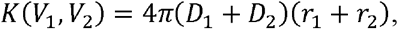

noting that 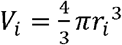 and that in the typical Stokes-Einstein case, *D*_*i*_ ∼ 1/*r*_*i*_ . We reasoned that in the fast quench and coalescence case, the simultaneously nucleating spheres were all of about the same size; for *r*_1_ = *r*_2_ , *K* is constant, and therefore the mergers were “equal opportunity”, giving an exponential size distribution. However, in the slow injection case, because the “older” spheres were much larger than newly injected spheres, *r*_1_ >> *r*_2_, and therefore *K*∼*r*_1_/*r*_2_, meaning that small spheres tended to merge with large spheres, allowing the “rich to get richer”, giving rise to a power law as observed in the slow injection case.

We therefore sought to test whether artificially strengthening the “rich get richer” effect by directly manipulating the kernel could give rise to a power-law size distribution starting from the same narrow initial size distribution as in our previous fast quench simulations, which had produced exponential distributions. Specifically, we performed simulations for ordinary diffusion *α* = 1, but for each simulation we set 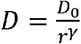 for different values of *γ*, where *γ* = 1 corresponds to the standard Stokes-Einstein dependence of the diffusion coefficient on radius. For values of *γ* < 1, merger events become biased towards larger spheres, and for *γ* < 0, spheres actually accelerate as they grow, strongly promoting preferential attachment (Fig. 6a).

**Fig. 6:**
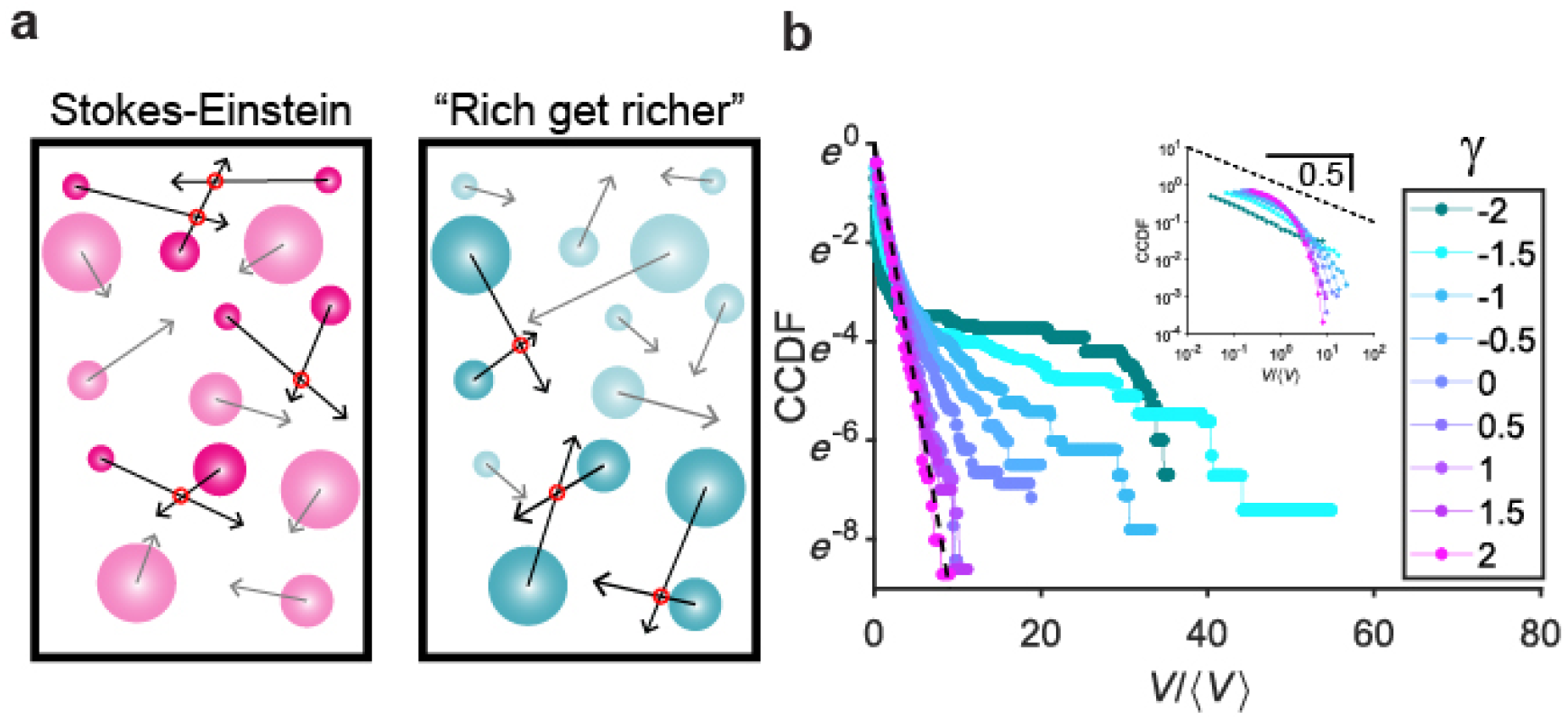
A “rich get richer” effect results in power-law-like size distributions. a. Schematic depicting the “rich get richer” effect. For the Stokes-Einstein diffusion of condensates, increased radius of large condensates is compensated for by a decrease in their mobility. Artificially inverting this relation results in increased collisions involving larger condensates, yielding a strong “rich get richer” effect. b. Cluster-size distributions were calculated varying the scaling exponent *γ* of the diffusion coefficient with sphere size (i.e., 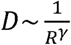) with *γ* ranging from -2 to 2 and were plotted as CCDFs in semi-log to compare to an exponential (dashed black line) in log-log to compare to a power law (inset).

We observed exponential distributions for *γ* > 1, consistent with the previous simulations, but as *γ* decreased below 1, we found progressively more power-law-like distributions (Fig. 6b). We then empirically calculated the coagulation kernel, estimating a collision rate constant by weighting each collision event by the number density of spheres of that size. We visualized the strength of the preferential attachment effect by plotting the value of the kernel diagonal against sphere size, verifying a far stronger preferential attachment effect in the *γ* = −2 simulations compared to the *γ* = 1 case (Fig. S6). We also tested several subdiffusive exponents with *γ* = 1, finding that the kernel diagonal remained constant as a function of radius for different subdiffusive exponents, consistent with the exponential distributions experimentally observed in the Corelet system. Thus, we were able to control the sphere-size distribution by mediating the strength of the “rich get richer” effect, independent of the rate of adding spheres to the system. We conclude that the scaling form of the size distribution generally reflects the strength of preferential attachment in the growth kinetics, whether due to material addition rate or to size-dependent diffusion, thus simply accounting for the scaling form of condensate size distributions in live-cell experiments in multiple systems.

## Discussion

In this work, we elucidated general principles underlying size distributions of intracellular condensates. We described two drastically different, experimentally motivated mechanisms of condensate growth: a quench-then-coalesce model, and a slow-nucleation model. We show that the size distributions of both can be accounted for by comparing the timescales of nucleation and coalescence, illuminating universal physical principles underlying vital biological processes.

The quench-then-coalesce mechanism of growth generically yields an exponential distribution, independent of subdiffusive exponent or time, which implies that a single parameter, i.e. the average condensate volume ⟨ *V*⟩, defines the distribution. The average condensate volume has previously been shown to grow in time as 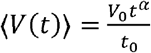 determined by the subdiffusive exponent *α*, following quench and nucleation to an average initial condensate volume *V*_*0*_ at some time *t*_0_ ^12^. We can therefore describe the distribution for the quench-then-coalesce mechanism simply as:

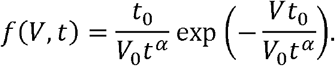

We note that this form is compatible with the self-similar dynamical scaling proposed by Vicsek and Family, who suggested in the context of diffusion-limited cluster aggregation that the distribution *f*(*s* ,*t)* of cluster size *s* at time *t*, can universally be described as 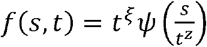 for some function *ψ*^39,40^.

Here, we have shown that both native nuclear speckles and synthetic condensates form via a quench-then-coalesce mechanism, which may present a common mechanism for size control of condensates. Indeed, several native organelles within the nucleus, including nuclear speckles^8^ but also nucleoli^23^, Cajal bodies^23^, histone locus bodies, paraspeckles, and PML bodies, all disassemble or scatter into the cytoplasm during mitosis^18^, likely due to a combination of dilution and post–translational modifications^19,41^, and then nucleate and regrow in the resulting daughter nuclei. Our results show that a relatively narrow, exponentially decaying size distribution will be attained for simple regrowth driven by coalescence. Furthermore, the growth of the mean condensate size will be slowed if the motion of condensates is subdiffusive, as is common in cells^20,42^, which may allow for the approximate maintenance of a characteristic mean condensate size over the duration of the cell cycle. Finally, while condensate nucleation often occurs at specific sites, e.g. at defined genomic loci, these loci themselves also undergo subdiffusive Brownian motion^31,43^. In general, we propose that the maintenance of an exponential size distribution of condensates by slow coalescence may be conveniently implemented given the natural dynamics of the cell cycle, irrespective of the molecular details of the condensate.

By contrast with the simple exponential distribution with a single mean size, power-law distributions are broad and scale-free, indicative in this context of a “rich get richer”, or preferential attachment effect, wherein coalescence events are biased towards larger, existing condensates. We demonstrated two possible causes of a preferential attachment phenomenon. In the case of the neurodegeneration-associated mutant Htt-polyQ species, we observed that ongoing nucleation effectively gives older condensates more time to grow, resulting in preferential attachment and a power-law distribution. There is not yet a consensus on whether small oligomers or larger Htt-PolyQ aggregates are to blame for accelerated disease progression or are a compensatory mechanism to slow cytotoxicity^44,45^. Regardless, the slow, steady production of aggregate generates a broad power-law distribution. We also suggest that a “rich get richer” effect is possible when diffusion is highly non-Stokesian. In general, the observation of power laws in condensates may be attributed to either or both of these mechanisms, or to others which have yet to be observed in cells, such as Ostwald ripening, which would effectively result in the merger of small and large condensates.

The distribution of condensate sizes is also relevant to other cellular pathologies, e.g., in nucleolar size changes in diseases such as progeria and cancer^9–11^. Here too, it is unclear whether the observed increase in condensate size in these contexts is causative or purely correlative. This may be indicative of altered coalescence or production dynamics, which might also be responsible for dysfunction. For instance, an exponential form with a larger mean might be indicative of an increased subdiffusive exponent, e.g. due to reduced nuclear elasticity which previously has been associated with cancer, while a broadened distribution would be indicative of a “rich-get-richer” effect. However, still other factors, such as nonequilibrium activity and condensate-dependent reaction rates^46^ have been described as potential regulators of condensate size. For example, nucleoli are commonly observed to exhibit an unexpected power-law distribution in size, which likely results from not only coalescence dynamics, but also from the strong influence of regulated steady-state production of ribosomal RNA. Examining such size distributions under perturbations that modulate biological activity represents an exciting next frontier, which will yield new insights into the coupling of classical materials coarsening dynamics and associated biological function and dysfunction.

## Methods

### Plasmids

eGFP-PolyQ74 was an adaptation of a gift from David Rubinsztein (Addgene plasmid #40262) with insertion of N-terminal residues MATLEK a part of the N17 domain and C-terminal residues PQAQPLLPQPQPPPPPPPPPPGPAVAEEPLHRP which comprise the PR (proline-rich) domain containing the C38 domain using primers as a part of In-Fusion cloning protocol (Takara). eGFP-PolyQ31 was adapted from eGFP-PolyQ74 through the variability of Q-length PCR products amplified using CloneAmp HiFi PCR Premix (Takara). FM5-NPM1-mCh was generated as described in ^27^.

To generate an sgRNA plasmid to target SRRM-2, a guide RNA duplex was designed to insert EYFP at the beginning of the endogenous human SRRM2 coding sequence, then cloned downstream the U6 promoter in the PX458 vector (Addgene Plasmid #48138; gift from Feng Zhang lab). The annealed oligonucleotides used to generate the final “pU6-SRRM2gRNA-CMV-FLAG-NLS-SpCas9-2A-mGFP” plasmid were as follows:

SRRM2 sgRNA Forward 5’ CACCGGGCCATGTACAACGGGATC 3’

SRRM2 sgRNA Reverse 5’ AAACGATCCCGTTGTACATGGCCC 3’

Separately, a gene fragment containing upstream and downstream SRRM2 homology arms, flanking the full-length EYFP coding sequence, was synthesized (IDT) then cloned into the pUC19 vector, first digested with HindIII and BamHI. The following oligonucleotides were used to create the final “pUC19-SRRM2 NonCode Homology-EYFP-SRRM2 Exon1 Homology” plasmid:

SRRM2 homology Forward 5’ TTGATGATAAGCTTcctttcttcaccactgagctccttcaaggg 3’

SRRM2 homology Reverse 5’ TTGATGATGGATCCccatagcctgcatgtccactcccagacgatgg 3’

Sanger sequencing (Genewiz) confirmed gene insertions in full for all constructs.

### Cell culture and cell line generation

U2OS (a kind gift from Mark Groudine lab, Fred Hutchinson Cancer Research Center), HeLa CCL-2 (ATCC), HEK-D (see below) and Lenti-X 293T (Takara) cells were cultured in growth medium consisting of Dulbecco’s modified Eagle’s medium (GIBCO), 10% fetal bovine serum (Atlanta Biologicals), and 10 U/mL Penicillin-Streptomycin (GIBCO), and incubated at 37°C and 5% CO_2_ in a humidified incubator.

#### iPSC culture

Induced pluripotent stem cells were obtained from Allen Institute for Cell Science at the Coriell Institute. The iPSC line AICS-0094-024 (Mono-Allelic mEGFP tagged SON WTC iPSC line) was used for our experiments. The colonies were expanded and maintained on Matrigel (Corning) in mTeSR Plus medium (Stem Cell Technologies). Cells were plated at 3000-10.000 cells per square centimeter in order to obtain ∼75% confluency every 5-7 days. The cells were passaged using ReLeSR (stem cell technologies) and split at a 1:10-1:50 ratio. mTeSR plus medium was supplemented with ROCK inhibitor Y-27632 (Selleckchem) for maximum 24 hours after cryopreservation or passaging. iPSCs were cryopreserved in mTeSR Plus medium supplemented with 30% Knock Out Serum Replacement (Gibco Life Technologies) and 10% DMSO.

#### CRISPR-Cas9 Based Generation of HEK293 Cells Expressing eYFP-tagged SRRM2

Single HEK293 cells (kind gift from Marc Diamond lab, UT Southwestern) were isolated by fluorescence activated cell sorting (FACS). A clonal line termed “HEK-D” was selected for its relatively flat morphology and preferable imaging characteristics. Cell line background was validated by STR profiling (America Type Culture collection).

To generate the EYFP-tagged SRRM2 clonal line, HEK-D cells were plated in a 24-well dish so at 70% confluency the next day. Lipofectamine-3000 (Thermo Fisher) was used to transfect the two plasmids at one-to-one ratio, according to manufacturer’s recommendation. Cells were amplified by successive passages to 6-well then 10-cm dishes. At ∼Day 10 following transfection, cells were trypsinized, pelleted, then re-suspended in flow cytometry buffer (DPBS with 10% fetal bovine serum). Single eYFP-positive cells were sorted by FACS (Flow Cytometry Resource Facility, Princeton Department of Molecular Biology) into separate wells of 96-well plates. Resulting colonies with expected localization of eYFP signal to nuclear speckles (but not cytoplasm) were amplified, and a single clone with relatively high fluorescence intensity was selected (“eYFP-SRRM2 48 HEK D”).

To validate, cells were passaged into single wells of a 96-well plate containing either 200 uL of pCRISPRv2-SRRM2 gRNA lentivirus (pooled, 6 gRNAs designed according to published recommendations^47^) or 200 uL of pCRISPRv2-NonTarget gRNA lentivirus. 72 hours later, confluent cells were washed, trypsinized, and passaged at 1:8 dilution factor. 96 hours later, confluent cells were plated onto a fibronectin-coated, glass-bottom, 96-well plate (MatTek), and eYFP fluorescence was compared between experimental (SRRM2 KO) and control (NonTarget) conditions, with confocal microscopy performed at 7-days post-lentivirus transduction. eYFP fluorescence was completely abrogated in the SRRM2 knockout (>90% of cells) condition, suggesting that eYFP was only integrated at endogenous SRRM2 loci.

Next, whole genome sequencing assessed the number of tagged copies in the hypotriploid HEK293 clone featuring three copies of chromosome 16, on which the SRRM2 gene is located. Two of the three copies contained the exogenous EYFP sequence between chr16:2,756,364 and chr16:2,756,365, corresponding to immediately prior to endogenous SRRM2’s start codon and after the expected upstream noncoding sequence. CRISPR-Cas9-mediated genome editing left a single nucleotide polymorphism encoding a synonymous codon (Gly6Gly) in tagged SRRM2 loci, otherwise unaltered from the parental sequence. Cells were then cultured and imaged as described elsewhere.

### Lentiviral transduction

For Corelet, Htt-PolyQ, and NPM-1 overexpression, lentivirus was produced by transfecting the transfer plasmids, pCMV-dR8.91, and pMD2.G (9:8:1, mass ratio) into LentiX cells grown to approximately 80% confluency in 6-well plates using FuGENE HD Transfection Reagent (Promega) per manufacturer’s protocol. A total of 3 μg plasmid and 9 μL of transfection reagent were delivered into each well. After 60-72 hours, supernatant containing viral particles was harvested and filtered with a 0.45 μm filter (Pall Life Sciences). Supernatant was immediately used for transduction or aliquoted and stored at −80°C. Cells were seeded at 10% confluency in 96-well plates and 20-200 μL of filtered viral supernatant was added to the cells. Media containing virus was replaced with fresh growth medium 24 hr post-infection. Infected cells were imaged no earlier than 72 hr after infection. For Htt-PolyQ HeLa lines, cells were transferred to a glass 96-well imaging plate coated with fibronectin two days following transduction and imaged 4-6 days post-transduction.

### Microscopy

Images of Corelets, nucleoli, and nuclear speckles were taken with a spinning-disk (Yokogawa CSU-X1) confocal microscope with a 100X oil immersion Apo TIRF objective (NA 1.49) and an Andor DU-897 EMCCD camera on a Nikon Eclipse Ti body. A 488nm laser for imaging GFP and global activation, and a 561 was used for imaging mCherry. The imaging chamber was maintained at 37°C and 5% CO_2_ (Okolab) with a 96 well plate adaptor. Z-stacks were taken every 300nm for 15 μm using an ASI MS-2000 stage controller. For drug treatments, DRB (Sigma) or ActD (Sigma) were dissolved in DMSO at 50mg/mL and 2mg/mL stock concentrations respectively, which was diluted in complete DMEM. Cells were treated by replacing media and incubating for 4 hours before imaging.

Images of Htt-PolyQ were taken on a Nikon A1 laser scanning confocal microscope using a 60x oil immersion lens with a numerical aperture of 1.4. A 488nm laser for imaging GFP. The imaging chamber was maintained at 37°C and 5% CO_2_ (Okolab) with a 96 well plate adaptor. Z-stacks were taken every 420nm for 15 μm using an ASI MS-2000 stage controller.

### Image Analysis

All images were analyzed in Fiji (ImageJ 1.52p)^48^ and MATLAB 2019b (Mathworks).

#### Corelets

Data was re-analyzed from previous work^20^. Individual nuclei were cropped by hand and saved as .tifs, which were analyzed in MATLAB. Droplets were segmented in the GFP channel by using Otsu’s algorithm to identify the nucleus in the first frame; an intensity threshold was defined as two standard deviations above the mean of the initial nuclear GFP intensity. This threshold was applied to all following frames to identify droplets; regions 4 pixels and smaller were discarded. Then, r*egionprops* was used to identify individual domains and calculate their area. Volumes were calculated by assuming perfect sphericity, based upon previous work ^20,49^; distributions were then calculated for each nucleus.

#### Nucleoli and nuclear speckles

Individual nuclei were cropped by hand and saved as .tifs, which were analyzed in MATLAB. Maximum projects were generated and used to calculate a threshold for each nucleus and channel, which was set to the mean of the image plus two standard deviations. This threshold was applied in three dimensions to each z-stack; *regionprops3* was used to calculate the volume of each condensate. Condensates below 20 voxels in size were discarded. CDFs were calculated for each nucleus, rescaled by mean volume, and averaged over nuclei. Error was propagated as standard error of the mean.

#### PolyQ aggregates

Individual cells were cropped by hand and saved as .tifs, which were analyzed in MATLAB. A frame of maximal mean cytoplasmic brightness was selected to calculate a threshold for the cytoplasm, which was set to the mean of the image plus 2.5 standard deviations. This threshold was applied in three dimensions to each z-stack; *regionprops3* was used to calculate the volume of each aggregate. Aggregates below 16 voxels in size were discarded. CDFs were calculated for each nucleus, rescaled by mean volume, and averaged over cellular cytoplasms. Error was propagated as standard error of the mean.

### Simulations

Simulations were performed in MATLAB on the Della cluster (Princeton Research Computing). First, an initial configuration of spheres were generated in 3D with periodic boundary conditions, with initial sphere volumes drawn from a uniform distribution ranging from 1 to 2 AU. Box size was set based on the volume fraction (generally set at 5%). Overlapping spheres were then merged. Sphere merger was implemented by randomly selecting a pair of overlapping spheres and replacing them with one new spheres centered at the center of mass of the pair. The size of the new sphere was determined by volume conservation of the original pair. This was iterated until no pairs of overlapping droplets remained. (For “vanishing” simulations employed as controls (Fig. S3,4) all spheres were set to a volume of 1 AU and maintained a volume of 1 AU after merger, resulting in spheres “vanishing”.) Then, for each sphere, a fractional Brownian motion trajectory with the appropriate α was synthesized (*wfbm*) for the appropriate number of timesteps (generally 10^5^). At each timepoint, each sphere proceeded one step along the synthesized trajectory, scaled such that *D*∼1/*r*^*γ*^. Spheres were merged as previously described. Merged spheres inherited the predetermined trajectory from one of their parent droplets (chosen arbitrarily), moving with the appropriate step size. This was repeated for the duration of the synthesized trajectories.

For injection simulations, the simulation box was empty initially; *J* spheres were initialized per timestep in randomly generated locations (i.e., if *J* = .25, a sphere was initialized every fourth timestep, while for *J* = 4, four spheres were initialized per timestep). Then, mergers and sphere diffusion proceeded as previously described before the next injection.

Size distributions were recorded for 20 replicates of each simulation condition, binned, and averaged before calculating CCDFs. To calculate the collision rate constant, 960 (20 in the case of the “vanishing” control simulations) simulation replicates were performed; for each merger event, the sphere size of the merging pair and number of spheres of that size were recorded. Each merger event occurring between spheres of volume *V*_1_ and *V*_2_ was weighted by a factor of 1/*f* (*V*_1_) *f* (*V*_2_), where *f* gives the number density of spheres of particular size. Summing over weights of all merger events in each simulation replicate and then averaging over replicates gave an estimate of the coagulation kernel value after normalization by system size and simulation duration (see Supplementary Note).

## Acknowledgements

This work was supported by NIH Grants U01 DA040601 and R01GM140032, the Howard Hughes Medical Institute, and the National Science Foundation, through the Center for the Physics of Biological Function (PHY-1734030) and the Graduate Research Fellowship Program (DCE-1656466, D.S.W.L.).

We thank Sofia Quinodoz for assistance with sequencing and cell line validation, members of the Wingreen and Brangwynne labs for useful discussion, Evangelos Gatzogiannis for microscopy assistance and the Princeton Flow Cytometry Resource Facility for FACS assistance.

## Author Declarations

### Conflict of Interest

C.P.B. is a founder and consultant for Nereid Therapeutics. All other authors declare no competing interests.

### Data and Reagent Availability

Data, reagents and software associated with this work are available upon reasonable request to the corresponding author(s).

