## Supplementary Notes and Figures for "A rich get richer effect governs intracellular condensate size distributions"

**Supplementary Information**

**Supplementary Note**

*Exponential distribution with size cutoff*

An exponential distribution is given by:

$f\left( x \right)=\frac{1}{\lambda}\exp(\frac{-x}{\lambda})$.

We consider an exponential distribution with a size cutoff at size $x_{0}$ :

$f\left( x \right)=\left\{ \begin{aligned} \begin{matrix} 0 & \mathrm{if} x<x_{0} \end{matrix} \\ \begin{matrix} \frac{N}{\lambda}\exp\left( \frac{-x}{\lambda} \right) & \mathrm{if} x>x_{0} \end{matrix} \end{aligned} \right.$,

for some normalization factor $N$. To calculate $N$*,* we realize that $\int_{0}^{\infty} f(x)=1$, and therefore $\int_{x_{0}}^{\infty} N(\frac{1}{\lambda}\exp\frac{-x}{\lambda})dx =1$, giving $N=\exp\left( \frac{x_{0}}{\lambda} \right)$.

Finally, to calculate the average,

$\langle x\rangle=\int_{x_{0}}^{\infty} \frac{1}{\lambda}\exp(\frac{-(x-x_{0})}{\lambda}) xdx=\lambda+x_{0}$,

and thus the mean is merely shifted, justifying a shift in the mean-variance plot corresponding to a minimum detection threshold (which can be applied computationally at a size scale greater than the diffraction limit to ensure proper sampling).

Higher central moments of the distribution are given in general by $\mu_{n}=\langle x-{\langle x\rangle}^{n}\rangle$, which should then be unchanged.

*Kernel Normalization*

The number of collisions $N$ between species of volumes $V_{1}$ and $V_{2}$ is given by

$$\frac{N(V_{1},V_{2})}{V_{\mathrm{sys}}}=\sum_{V_{1},V_{2}} \int_{t} K(V_{1},V_{2})f(V_{1},t)f(V_{2},t)dt$$

Then $I(V_{1},V_{2})$ empirically counts collisions between populations of volumes $V_{1}$ and $V_{2}$*,* so

$$I(V_{1},V_{2})=K(V_{1},V_{2})V_{\mathrm{sys}}f(V_{1},t)f(V_{2},t)\int dt$$

And thus the number of collisions counted over a period of time $T$ in a system of volume size $V_{\mathrm{sys}}$ is given by:

$K(V_{1},V_{2})=\frac{I(V_{1},V_{2})}{f(V_{1},t)f(V_{2},t)V_{\mathrm{sys}}T}$.

**Supplementary Figures**

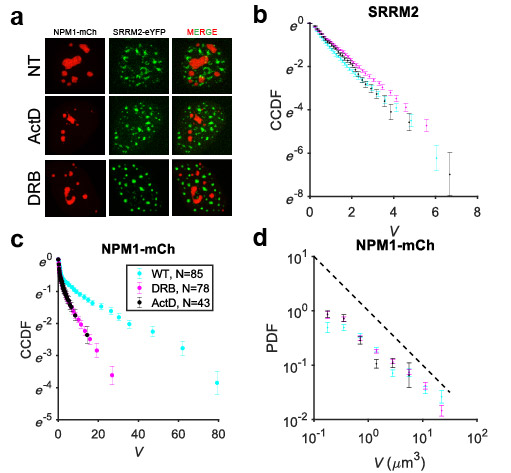

**Fig. S1 Analysis of endogenous nuclear body size distributions upon transcriptional inhibition.**

a. HEK-D cells were endogenously tagged with SRRM-2 to label nuclear speckles and were lentivirally induced to express exogenous NPM1-mCh to label nucleoli. Cells were treated with either 50μg/mL DRB for 4 hours, or with 10μg/mL actinomycin-D for 2 hours. Example nuclei are shown for each condition and channel.
b. Nuclear speckles demonstrated exponential distributions as seen by plotting the CCDF for all three drug conditions; ActD decreased speckle size to $1.05\pm0.03 \mu m^{3}$ (mean$\pm$SEM, $N=43$ nuclei) compared to nontreated control ($\sim1.16 \mu m^{3}$), but less than DRB ($\sim.92 \mu m^{3})$.
c. Unlike speckles, nucleoli strongly deviate from an exponential, exhibiting nonlinear behavior in the semi-log CCDF. Nucleoli treated with transcriptional inhibitors were also much smaller than the nontreated control, whose average volume was $11.6\pm0.8 \mu m^{3}$($N=85$). The average volume after treatment with DRB was $4.0\pm0.2 \mu m^{3}$ ($N=78$), and $0.97\pm.02 \mu m^{3}$ after treatment with ActD ($N=43$).
d. Binning and averaging demonstrates that a power law with a slope slightly less than 1 (black dashed reference line) was exhibited over a limited range (approximately 1-1.5 decades) for nucleoli, consistent with literature.

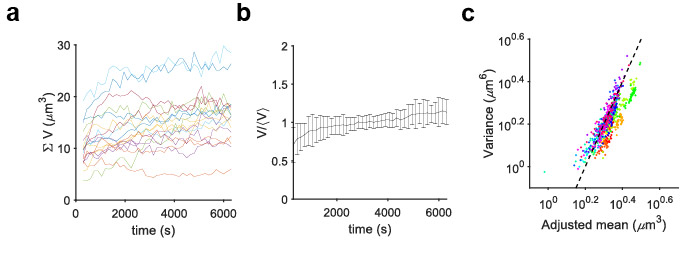
**Fig. S2 The Corelet system maintains a relatively constant amount of dense phase material and closely follows an exponential distribution.**

a. For 18 nuclei, the total volume of Corelet condensates in each nucleus was calculated and plotted over time (imaging every 3 seconds following the first 5 minutes of activation).
b. To visualize overall changes in total condensate volume, in each of the 18 nuclei the total Corelet condensate volume was normalized by the time-averaged total condensate volume and then averaged over nuclei. Error bars reflect standard deviation.
c. It was noted that the mean in Fig. 2D is horizontally shifted from the expectation line; however, for an exponential distribution a shift in the mean can be attributed to a minimum experimentally detectable size, e.g., due to the diffraction limit (see Supplementary Note). The mean condensate size was therefore adjusted by adding the minimum detection threshold corresponding to a condensate of radius $0.3 \mu m$ and replotted with respect to variance on a log-log plot for comparison to the expectation from an exponential distribution (black dashed line).

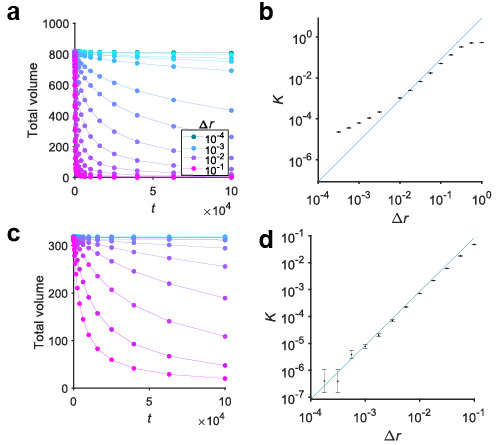

**Fig. S3 Control simulations validate merger rates in coalescence simulations.**

a. To more accurately obtain the merger rate of spheres in our simulation framework, we first modified the simulation such that spheres do not change in size upon merger, effectively lowering the amount of total material in the system over time but providing a more accurate estimate of the collision rate constant $K$, since all spheres are of the same size. We performed this simulation beginning with a system of 1000 spheres each of volume 1; spheres overlapping in the initial condition were replaced by a single sphere, and the system was allowed to evolve according to the specified diffusive step size. Total volume was plotted after the initial condition was resolved.
b. The theoretically predicted merger rate (blue line; see Supplementary Note) was compared to the input diffusive step size for a range of conditions (black points), finding agreement over a limited range of step-size values. Error bars reflect standard error of the mean over 20 replicates.
c. To determine the source of disagreement between theoretical prediction and apparent merger rate in (b), we first performed a “pre-run” of $2\times{10}^{4}$ timesteps at a step size $\Delta x=0.03$ starting with an initial condition identical to that in (a) for all conditions before collecting data, effectively “diluting” the system by nearly 5-fold.
d. The collision rate constant was calculated for a range of step-size values following the pre-run protocol, revealing a closer agreement with the theoretical prediction over a larger range of values.

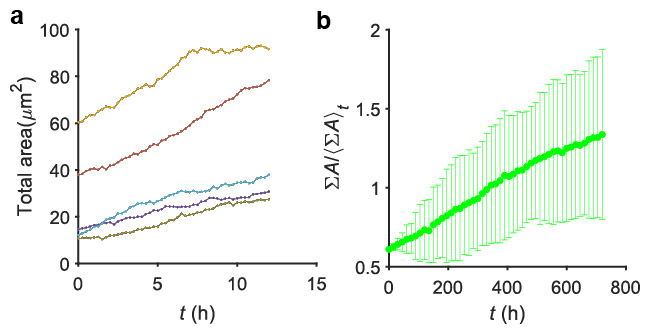
**Fig. S4 PolyQ dynamics reveal a large increase in aggregate material over time.**

a. Total PolyQ protein aggresome area was plotted over time for each cell, imaging every 15 minutes for 12 hours.
b. To assess the overall change in aggregate material, the aggregate area normalized by the time averaged aggregate area was averaged over cells and plotted; error bar reflects standard deviation. The amount of aggregate material increases on average about twofold from the beginning to the end of the time course.

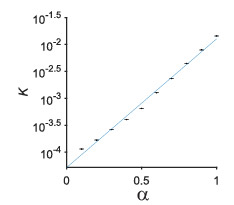

**Fig. S5 Empirical calculation of collision rate constant dependence on subdiffusive coefficient**

a. The rate constant dependence on $\alpha$ was empirically calculated by running 20 replicates of simulations containing 1000 spheres which did not change size upon merger (i.e., identical to Fig. S2a), with $\Delta r=.05$ and varying $\alpha$ from 0.1 to 1. $K_{\alpha}$ was calculated for each condition by taking the merger probability weighted by the sphere number density. Error bar reflects standard error of the mean over replicates. The $K_{\alpha}$ value was well fit by a linear model ($R^{2}=0.993)$ in semi-log, i.e., $\log(K_{\alpha}) \sim\alpha$ (blue line), which was used to calculate $\chi$ (Fig. 4).

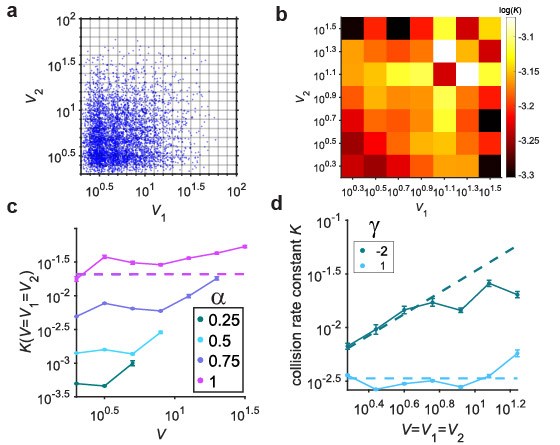

**Fig. S6 Calculating the coagulation kernel allows for quantification of the “rich get richer” effect.**

a. To characterize the simulated merger dynamics, we first plotted the incidence of individual merger events between spheres of particular volumes for an example system (5000 randomly chosen events were chosen to plot for visual clarity).
b. To describe the apparent strength of the preferential attachment effect, we weighted each merger event by the number density of the sphere sizes involved in the merger, and averaged over each possible pair of sizes, giving an estimate of the collision rate constant or coagulation kernel.
c. The coagulation kernel diagonal was computed for 960 replicates of the fast quench and coalescence simulation with 1000 spheres and plotted against theoretical expectation for ordinary diffusion (magenta dashed line), demonstrating that for all subdiffusive conditions, there is no strong preferential attachment effect, as expected.
d. The diagonal of the kernel (e.g., as shown in Fig. 5b) was calculated and plotted for simulations with $\gamma=1$ and $\gamma=-2$ for 1000 replicates. Both conditions were compared to and agreed well with theoretical expectation, i.e., $K\left( V \right)=16\pi{\Delta x^{2}V}^{\frac{1-\gamma}{3}}$ (dashed lines).
